# Model-based fMRI reveals co-existing specific and generalized concept representations

**DOI:** 10.1101/2020.05.26.117507

**Authors:** Caitlin R. Bowman, Takako Iwashita, Dagmar Zeithamova

## Abstract

There has been a long-standing debate about whether categories are represented by individual category members (exemplars) or by the central tendency abstracted from individual members (prototypes). Across neuroimaging studies, there has been neural evidence for either exemplar representations or prototype representations, but not both. In the present study, we asked whether it is possible for individuals to form multiple types of category representations within a single task. We designed a categorization task to promote both exemplar and prototype representations, and we tracked their formation across learning. We found evidence for co-existing prototype and exemplar representations in brain in regions that aligned with previous studies: prototypes in ventromedial prefrontal cortex and anterior hippocampus and exemplars in inferior frontal gyrus and lateral parietal cortex. These findings show that, under the right circumstances, individuals may form representations at multiple levels of specificity, potentially facilitating a broad range of future memory-based decisions.

---

The ability to form new conceptual knowledge is a key aspect of healthy memory function. There has been a longstanding debate about the nature of the representations underlying conceptual knowledge, which is exemplified in the domain of categorization. Some propose that categories are represented by their individual category members and that generalizing the category label to new examples involves joint retrieval and consideration of individual examples encountered in the past (i.e., exemplar models, Figure 1A; Kruschke, 1992; Medin, Schaffer, & College, 1978; Nosofsky, 1986). Others propose that categories are represented by their central tendency – an abstract prototype containing all the most typical features of the category (i.e., prototype models, Figure 1B; Homa, Cross, Cornell, Goldman, & Shwartz, 1973; Posner & Keele, 1968; Reed, 1972). Category generalization then involves consideration of a new item’s similarity to relevant category prototypes.

**Figure 1.**
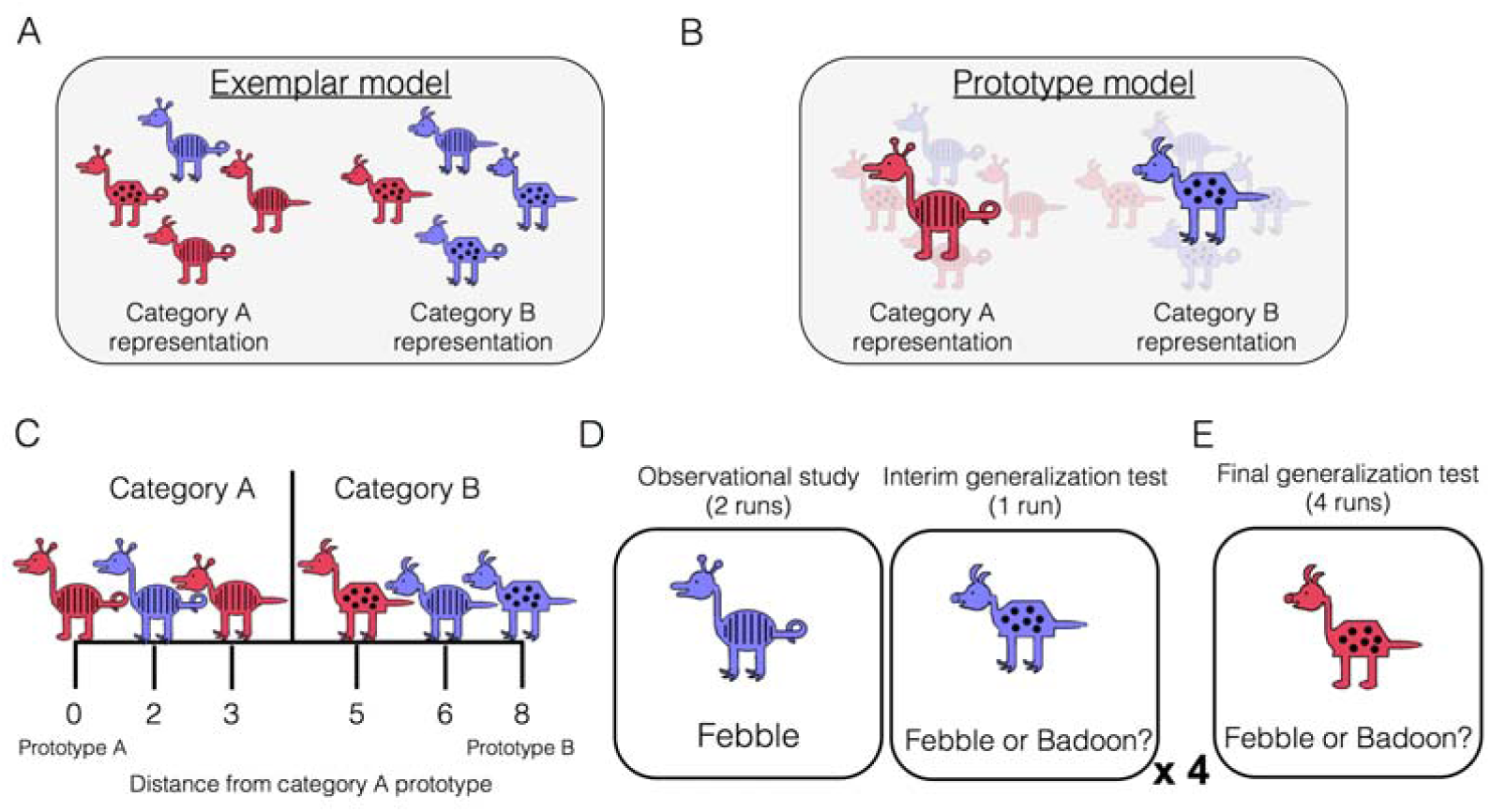
Category-learning task. Conceptual depiction of **A**. exemplar and **B**. prototype models. Exemplar: categories are represented as individual exemplars. New items are classified into the category with the most similar exemplars. Prototype: categories are represented by their central tendencies (prototypes). New items are classified into the category with the most similar prototype. **C**. Example stimuli. The leftmost stimulus is the prototype of category A and the rightmost stimulus is the prototype of category B, which shares no features with prototype A. Members of category A share more features with prototype A than prototype B, and vice versa. **D**. During the learning phase, participants completed four study-test cycles while undergoing fMRI. In each cycle, there were two runs of observational study followed by one run of an interim generalization test. During observational study runs, participants saw training examples with their species labels without making any responses. During interim test runs, participants classified training items as well as new items at varying distances. **E**. After all study-test cycles were complete, participants completed a final generalization test that was divided across four runs. Participants classified training items as well as new items at varying distances.

Both the prototype and exemplar accounts have been formalized as quantitative models and fit to behavioral data for decades, with numerous studies supporting each model (exemplar meta-analysis: Nosofsky, 1988; prototype meta-analysis: Smith & Minda, 2000). Neuroimaging studies have also provided support for these models. Studies using univariate contrasts showed overlap between neural systems supporting categorization and recognition (Nosofsky, Little, & James, 2012), as well as medial temporal lobe involvement in categorization (Koenig et al., 2008; Nomura et al., 2007), both of which have been interpreted as indicating a role of exemplar retrieval in categorization. More recently, studies have used parameters generated from formal prototype and exemplar models with neuroimaging data, but with conflicting results. Mack and colleagues (2013) found similar behavioral fits for the two models, but much better fit of the exemplar model to brain data. Parts of the lateral occipital, lateral prefrontal and lateral parietal cortices tracked exemplar model predictors while no region tracked prototype predictors. The authors concluded that categorization decisions are based on memory for individual items rather than abstract prototypes. In contrast, Bowman and Zeithamova (2018) found better fit of the prototype model in both brain and behavior. The ventromedial prefrontal cortex and anterior hippocampus tracked prototype predictors, consistent with their role in episodic inference through memory integration of related episodes (Schlichting, Mumford, & Preston, 2015; Shohamy & Wagner, 2008; Zeithamova, Dominick, & Preston, 2012). Thus, although the hippocampus is typically thought to support memory for specific episodes, its role in categorization goes beyond retrieval of specific instances to include the formation of generalized prototype representations.

It is possible that the seemingly conflicting findings regarding the nature of category representations arose because individuals are capable of forming either type of representation. Prior studies have compared different category structures and task instructions to identify multiple memory systems supporting categorization (e.g., Aizenstein et al., 2000; Ashby, Alfonso-Reese, Turken, & Waldron, 1998; Ell, Weinstein, & Ivry, 2010; Zeithamova, Maddox, & Schnyer, 2008). While such findings show that the nature of concept representations depend on task demands, it is unclear if both prototype and exemplar representations can co-exist within the same task. To test this idea, we used fMRI in conjunction with a categorization task designed to balance encoding of individual examples vs. abstract information. This task used a training set with examples relatively close to the prototype, which has been shown to promote prototype abstraction (Bowman & Zeithamova, 2018, in press). To promote exemplar encoding, we used an observational training task rather than feedback-based training (Cincotta & Seger, 2007; Heindel, Festa, Ott, Landy, & Salmon, 2013; Poldrack et al., 2001). We then measured the extent to which both prototype- and exemplar-tracking regions could be identified, focusing on the VMPFC and anterior hippocampus as predicted prototype-tracking regions, and lateral occipital, prefrontal, and parietal regions as predicted exemplar-tracking regions.

We also asked whether there are shifts across learning in the type of concept representation individuals rely on to make categorization judgments. While some have suggested that memory systems compete with one another during learning (Poldrack & Packard, 2003; Seger & Cincotta, 2005), prior studies fitting exemplar and prototype models to fMRI data have done so only during a categorization test that followed extensive training, potentially missing dynamics occurring earlier in concept formation. Notably, memory consolidation research suggests that memories become abstract over time, often at the expense of memory for specific details (McClelland, McNaughton, & O’Reilly, 1995; Moscovitch, Cabeza, Winocur, & Nadel, 2016; Payne et al., 2009; Posner & Keele, 1970), suggesting that early concept representations may be exemplar-based. In contrast, research on schema-based memory shows that abstract knowledge facilitates learning of individual items by providing an organizational structure into which new information can be incorporated (Bransford & Johnson, 1972; Tse et al., 2007; van Kesteren, Ruiter, Fernández, & Henson, 2012). Thus, early learning may instead emphasize formation of prototype representations, with exemplars emerging later. Finally, abstract and specific representations need not trade-off in either direction. Instead, the brain may form these representations in parallel (Collin, Milivojevic, & Doeller, 2015; Schlichting et al., 2015), generating the prediction that both prototype and exemplar representations may grow in strength over the course of learning.

In the present study, participants underwent fMRI scanning while learning two novel categories or ‘species,’ which were represented by cartoon animals varying on eight binary dimensions (Figure 1C). The learning phase consisted of two types of runs: observational study runs and interim generalization test runs (Figure 1D). During study runs, participants passively viewed individual category members with their accompanying species label (‘Febble’ or ‘Badoon’). All of the items presented during study runs differed by two features from their respective prototypes (for example, exemplars depicted in Figure 1A). After completing two runs of observational study, participants underwent an interim generalization test run in which participants classified cartoon animals into the two species. Test items included the training items as well as new items at varying distances from category prototypes. Across the entire learning phase, there were 4 study-test cycles, with different new test items at every cycle. The learning phase was followed by a final generalization test, whose structure was similar to the interim test runs but more extensive (Figure 1E). To test for co-existing prototype and exemplar correlates in the brain during interim and final generalization tests, we used latent metrics generated from each model as trial-by-trial predictors of BOLD activation in six regions of interest (Figure 2): ventromedial prefrontal cortex, anterior hippocampus, posterior hippocampus, lateral occipital cortex, inferior frontal gyrus, and lateral parietal cortex. To identify potential changes with learning, we tested these effects separately in the first half of the learning phase (interim tests 1 and 2) and second half of the learning phase (interim tests 3 and 4) as well as in the final test.

**Figure 2.**
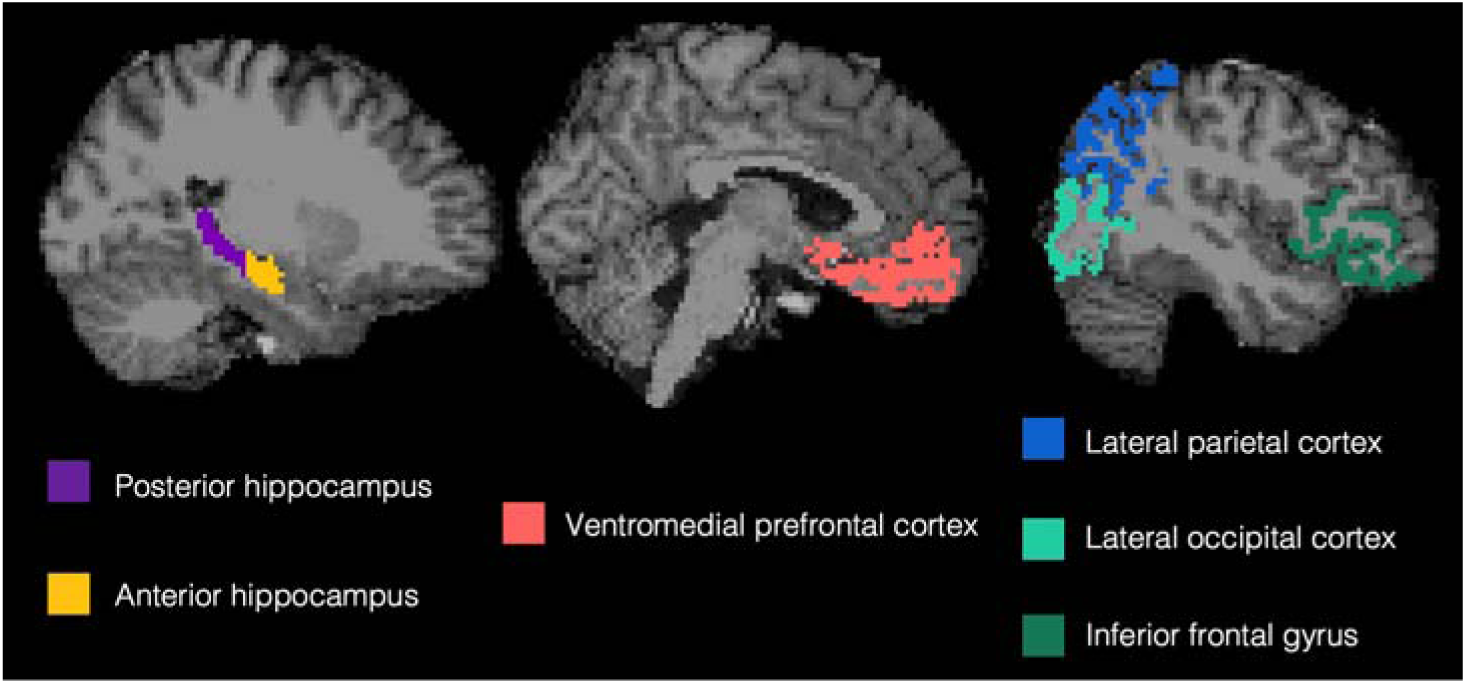
Regions of interest from a representative subject. Regions were defined in the native space of each subject using automated segmentation in Freesurfer.

As a complementary approach, we used pattern similarity analyses on the observational study runs to identify regions representing item-level information (i.e., individual category members) and/or category-level information. Item-level information was defined as greater neural pattern similarity for repetitions of a specific stimulus compared to its neural similarity to other stimuli. Category-level information was defined as greater pattern similarity for items in the same category compared to physically equally similar items from different categories. We were interested whether such a distinct analysis method, applied on a distinct portion of neuroimaging data (observational learning rather than categorization task), would provide converging or complementary information to model-based MRI results. As with neural prototype and exemplar model fits, we compared the first and second half of learning to identify potential representational shifts.

## Results

### Behavioral

#### Accuracy

##### Interim tests

Categorization performance across the four interim tests is presented in Figure 3A. We first tested whether generalization accuracy improved across the learning phase and whether generalization of category labels to new items differed across items of varying distance to category prototypes. There was a significant main effect of interim test number [*F*(3,84) = 3.27, *p* =.03, 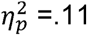], with a significant linear effect [*F*(1,28) = 9.91, *p* =.004, 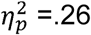] driven by increasing generalization accuracy across the interim tests. There was also a significant main effect of item distance [*F*(3,84) = 51.75, *p* < .001, 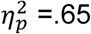] with a significant linear effect [*F*(1,28) = 126.04, *p* < .001, 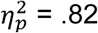] driven by better accuracy for items closer to category prototypes. The interim test number x item distance interaction effect was not significant [*F*(9,252) = 0.62, *p* =.78, 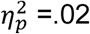]. We next tested whether accuracy for old training items was higher than new items of the same distance (i.e., distance 2) and whether that differed over the course of the learning phase. There was a linear effect of interim test number [*F*(1,28) = 16.78, *p* < .001, 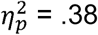] driven by increasing accuracy across the tests. There was also a significant main effect of item type (old vs. new) [*F*(1,28) = 8.76, *p* =.01, 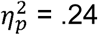], driven by higher accuracy for old items (M = .83, SD =.11) relative to new items of the same distance from the prototypes (M = .77, SD = .10). The interim test number x item type interaction effect was not significant [*F*(3,84) = 0.35, *p* = .79, 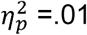], indicating that the advantage for old compared to new items was relatively stable across learning.

**Figure 3.**
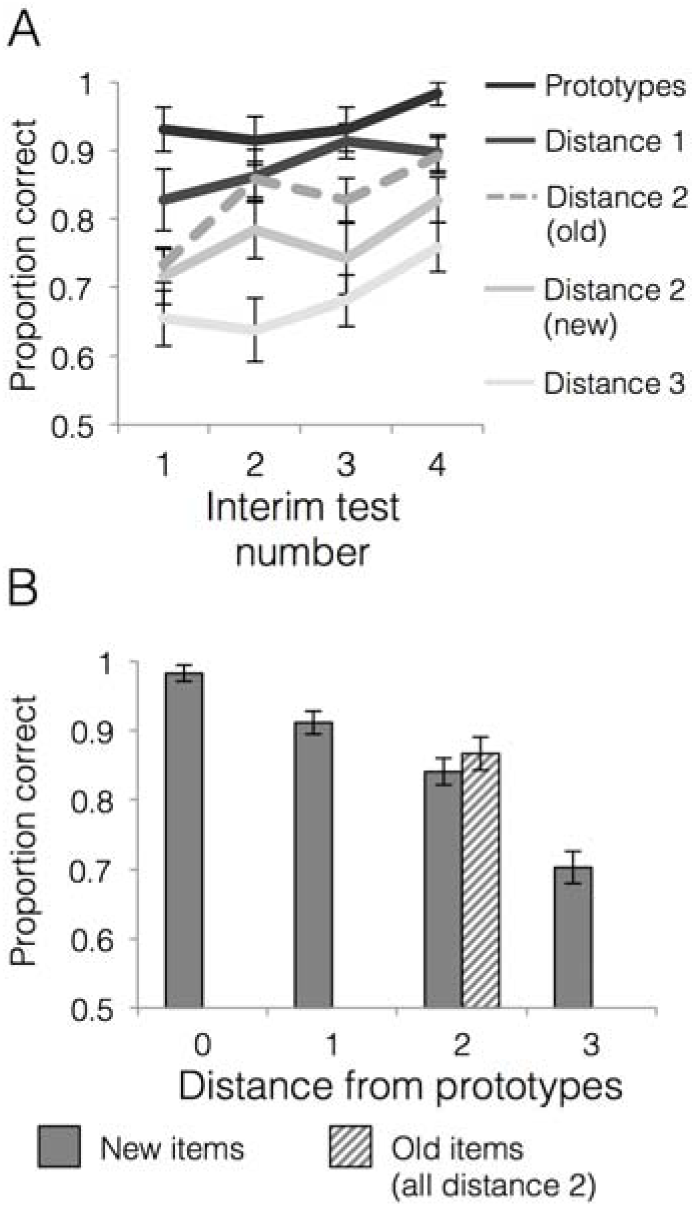
Behavioral accuracy for interim and final tests. **A**. Mean generalization accuracy across each of four interim tests completed during the learning phase. **B**. Mean categorization accuracy in the final test. In both cases, accuracies are separated by distance from category prototypes (0-3) and old vs. new (applicable to distance 2 items only). Error bars represent the standard error of the mean.

##### Final test

Accuracies for generalization items at each distance from the prototype as well as for training items (all training items were at distance 2 from the prototypes) are presented in Figure 3B. A repeated measures ANOVA on new items that tested the effect of distance from category prototypes on generalization accuracy showed a main effect of item distance [*F*(3,84) = 53.61, *p* < .001, 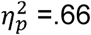] that was well characterized by a linear effect [*F*(1,28) = 124.55, *p* <.001, 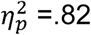]. Thus, the categorization gradient driven by higher accuracy for items closer to category prototypes observed during learning was also strong during the final test. In contrast, a paired t-test for accuracy on old relative to new items at distance two showed that the numeric advantage for old relative to new items was not statistically significant in the final test [*t*(28) = 0.93, *p* = .36, *CI*_95_[-.03,.08], *d* = 0.22].

#### Behavioral Model fits

Figure 4 presents model fits in terms of raw negative log likelihood for each phase (lower numbers mean lower model fit error and thus better fit). We first compared model fits for interim tests across the first and second half of the learning phase. There was a significant main effect of learning phase [*F*(1,28) = 39.74, *p* <.001, 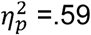] with better model fits (i.e., lower error) in the second half of the learning phase (M = 5.98, SD = 5.81) compared to the first half (M = 10.64, SD = 6.72). There was also a significant main effect of model [*F*(1,28) = 17.50, *p* <.001, 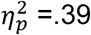] with better fit for the prototype model (M = 7.86, SD = 5.95) compared to the exemplar model (M = 8.77, SD = 6.02). The learning phase x model interaction effect was not significant [*F*(1,28) = 0.01, *p* =.91, 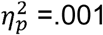], with a similar prototype advantage in the first half (d = 0.13, *CI*_95_[0.31,1.45]) as in the second half (d = 0.16, *CI*_95_[0.22,1.65]). When we compared prototype and exemplar model fits in the final test, we again found a significant advantage for the prototype model over the exemplar model [*t*(28) = 3.53, *p* =.001, *CI*_95_[0.89, 3.39], *d* = 0.23].

**Figure 4.**
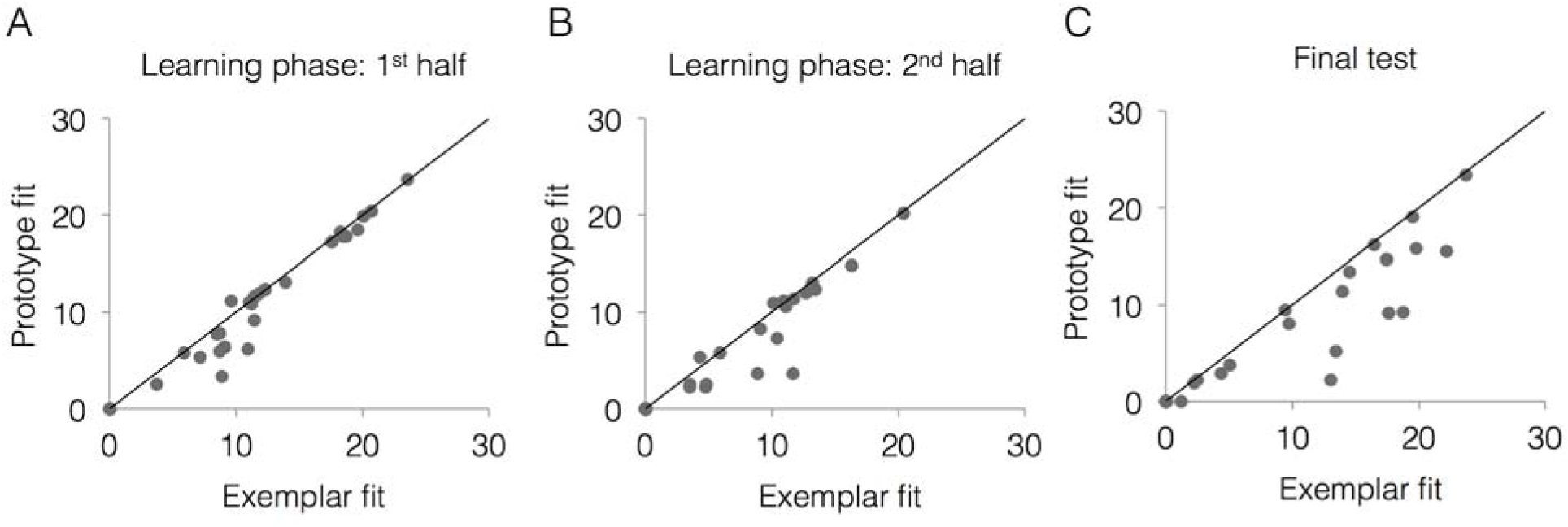
Relative exemplar vs. prototype model fits for each subject. **A**. The 1^st^ half of the learning phase (interim tests 1-2). **B**. The 2^nd^ half of the learning phase (interim tests 3-4). **C**. The final test. Fits are given in terms of negative log likelihood (i.e., model error) such that lower values reflect better model fit. Each dot represents a single subject and the trendline represents equal prototype and exemplar fit. Dots above the line have better exemplar relative to prototype model fit. Dots below the line have better prototype relative to exemplar model fit.

Thus, the prototype model provided an overall better fit to behavioral responses throughout the learning phase and final test, and the effect size of the prototype advantage was largest in the final test.

### fMRI

#### Model-based MRI

##### Learning phase

We first tested the degree to which prototype and exemplar information was represented across different brain regions and across different points of the learning phase. Using the data from the interim generalization tests, we compared neural model fits across our six ROIs across the first and second half of the learning phase. Full ANOVA results are presented in Table 1. Figure 5 presents neural model fits for each ROI. Figure 5A represents 1^st^ half of the learning phase, Figure 5B represents the 2^nd^ half of the learning phase, and Figure 5C represents fits collapsed across the entire learning phase (to illustrate the main effects of ROI, model and ROI x model interaction).

**Table 1.** ANOVA results for model-based fMRI during the learning phase.

| Table 1. ANOVA results for model-based fMRI during the learning phase |  |  |  |  |
| --- | --- | --- | --- | --- |
| Effect | df | F | p | $\eta_p^2$ |
| ROI | 3.4,95.6 GG | 3.90 | .002 | .12 |
| Model | 1,28 | 2.60 | .12 | .09 |
| Learning half | 1,28 | 2.18 | .15 | .07 |
| ROI x Model | 2.9,80.3 GG | 5.91 | .001 | .17 |
| ROI x Learning half | 3.1,86.9 GG | 0.53 | .67 | .02 |
| Model x Learning half | 1,28 | 0.09 | .76 | .003 |
| ROI x Model x Learning Half | 3.2,89.6 GG | 2.31 | .08 | .08 |

**Figure 5.**
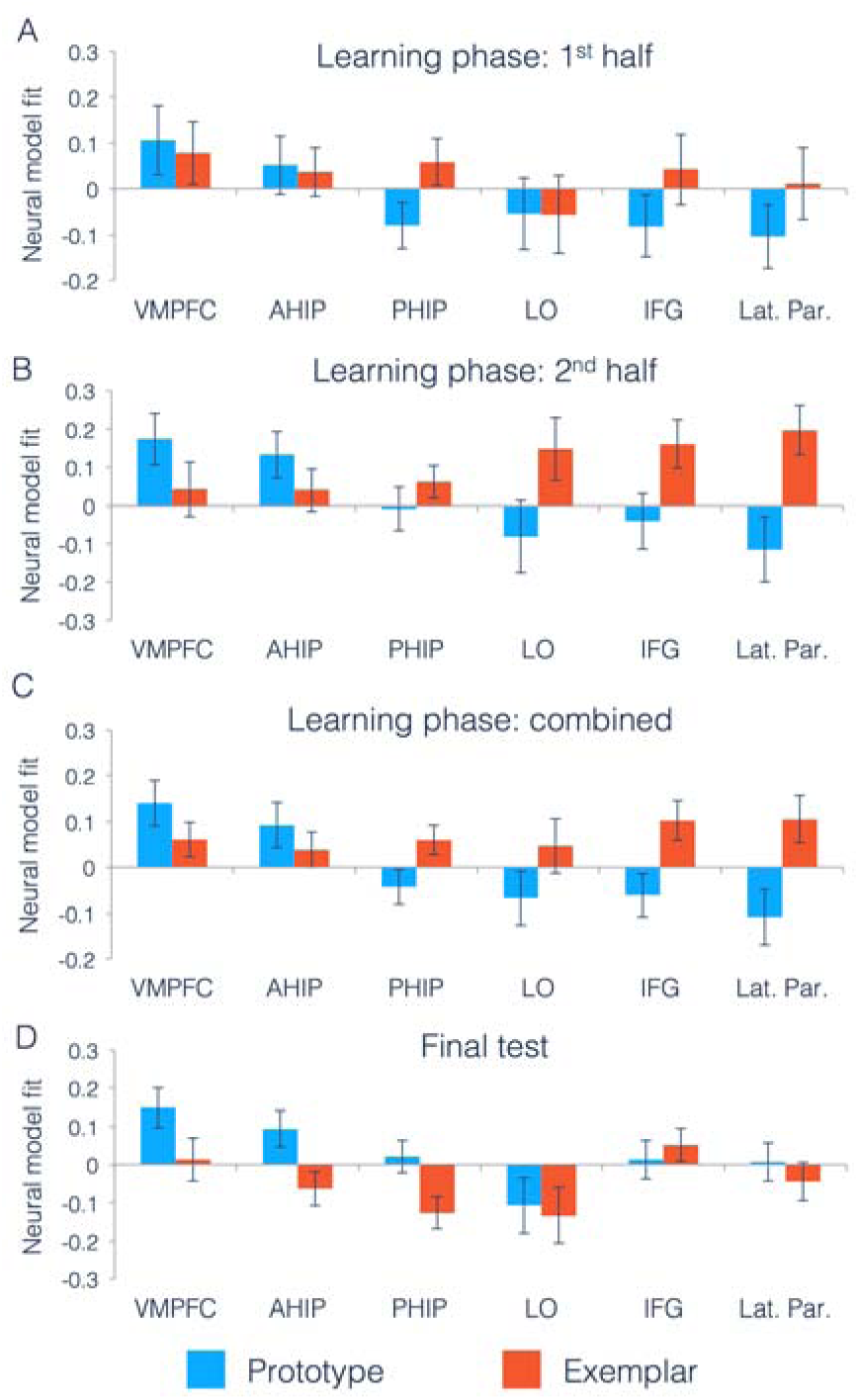
Neural prototype and exemplar model fits. Neural model fits for each region of interest for **A**. the first half of the learning phase, **B**. the second half of the learning phase, **C**. the overall learning phase (averaged across the first and second half of learning), and **D**. the final test. Prototype fits are in blue, exemplar fits in red. Neural model fit is the effect size: the mean/SD of ß-values within each ROI, averaged across appropriate runs. VMPFC = ventromedial prefrontal cortex, ahip = anterior hippocampus, phip = posterior hippocampus, LO = lateral occipital cortex, IFG = inferior frontal gyrus, and Lat. Par. = lateral parietal cortex.

As predicted, there was a significant ROI x Model interaction effect, indicating that there were differences across regions in the type of category information that they tracked. To understand the nature of this interaction, we computed follow-up t-tests on the neural model fits in each ROI, collapsed across the first and second half of the learning phase. Consistent with prior work (Bowman & Zeithamova, 2018), the VMPFC and anterior hippocampus (our predicted prototype regions) significantly tracked prototype information [VMPFC: *t*(28) = 2.86, *p* =.004, *CI*_95_[µ > .06], *d* = 0.75; anterior hippocampus: *t*(28) = 1.88, *p* =.04, *CI*_95_[µ > 0.009], *d* = 0.49]. For the predicted exemplar regions, we found that both lateral parietal cortex and inferior frontal gyrus significantly tracked exemplar model predictions [lateral parietal: *t*(28) = 2.06, *p* = 0.02, *CI*_95_[µ > 0.02], *d* = 0.54; inferior frontal: *t*(28) = 2.40, *p* = 0.01, *CI*_95_[µ > 0.03], *d* = 0.63], with numerically positive exemplar correlates in lateral occipital cortex that were not statistically significant [*t*(28) = 0.78, *p* = 0.22, *CI*_95_[µ > -0.05], *d* = 0.20].

As in our prior study, the posterior hippocampus showed numerically better fit of the exemplar predictor, but neither the exemplar effect [*t*(28) = 1.88, p =.07, *CI*_95_[-.01,.13], *d* = 0.49] nor the prototype effect reached significance [*t*(28) = -1.14, *p* =.26, *CI*_95_[-.12,.03], *d* = 0.30]. Comparing the effects in the two hippocampal regions as part of a 2 (hippocampal ROI: anterior, posterior) x 2 (model: prototype, exemplar) repeated-measures ANOVA, we found a significant interaction [*F*(1,28) = 9.04, *p* = .006, 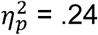], showing that there is a dissociation along the hippocampal long axis in the type of category information represented. Taken together, we found evidence for different types of category information represented across distinct regions of the brain.

We were also interested in whether there was a shift in representations that could be detected across learning. The only effect that included learning phase that approached significance was the three-way ROI x model x learning phase interaction, likely reflecting the more pronounced region x model differences later in learning (Figure 5A vs. Figure 5B).

##### Final test

Figure 5D presents neural model fits from each ROI during the final test. We tested whether the differences across ROIs identified during the learning phase were also present in the final test. As during the learning phase, we found a significant main effect of ROI [*F*(2.9,79.8) = 9.13, *p* <.001, 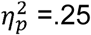, GG] and no main effect of model [*F*(1,28) = 1.65, *p* = .21, 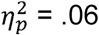]. However, unlike the learning phase, we did not find a significant model x ROI interaction effect [*F*(3.3,91.2) = 1.81, *p* = .15, 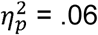, GG]. Because this was a surprising finding, we wanted to better understand what had changed from the learning phase to the final test. Thus, although the ROI x model interaction was not significant in the final test, we computed follow-up tests on regions that had significantly tracked prototype and exemplar predictors during the learning phase. As in the learning phase, both the VMPFC and anterior hippocampus continued to significantly track prototype predictors during the final test with effect sizes similar to those observed during learning [VMPFC: *t*(28) = 2.83, *p* =.004, *CI*_95_[µ > 0.06], *d* = 0.74; anterior hippocampus: *t*(28) = 1.98, *p* = .03, *CI*_95_[µ >0.01], *d* = 0.52]. However, exemplar correlates did not reach significance in any of the predicted exemplar regions (all t < 1.18, p > .12, d < 0.31).

#### Pattern similarity

As a complement to model-based fMRI, we used pattern similarity analyses during observational study runs to measure item-level and category-level information. Observational study runs were well suited to pattern similarity analyses because, unlike test runs, there was no required motor response, and the physical similarity of training items was well controlled, allowing us to detect category effects that were not driven by differences in the number of shared features for pairs of items within the same category vs. between categories. First, we computed neural pattern similarity between all pairs of trials from different runs (e.g., trial 1 from run 1 to trial 1 from run 2, trial 1 from run 1 to trial 2 from run 2, etc.) to avoid any possible issues with within-run autocorrelation (Mumford, Davis, & Poldrack, 2014). We then sorted the neural pattern similarity values into three bins, depending on whether they reflected the similarity between activation patterns for repetitions of the same training item, two separate training items that belonged to the same category, or two training items that belonged to different categories. We then computed item-level representations as the difference between pattern similarity for same item and pattern similarity for distinct items from the same category. Category-level representations were defined as the difference between same category similarity and different category similarity. As all items in the same category differed from one another by four features, we limited the different category pattern similarity analysis to only those pairs of items that also differed from one another by four features, which was by design the vast majority of all possible different category item pairs. This allowed us to measure category representations that were not merely driven by differences in physical similarity as all similarity comparisons were based on items that differed by four features.

Pattern similarity results are presented separately for each ROI and each learning half for category-level information (Figure 6A) and item-level information (Figure 6B). A 6 (ROI) x 2 (learning half) repeated-measures ANOVA for category-level representations did not reveal any significant effects [main effect of ROI: *F*(2.5,68.9) = 0.70, *p* = .62, 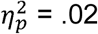, GG; main effect of learning half: *F*(1,28) = 0.10, *p* = .75, 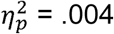; ROI x learning half interaction: *F*(3.5,97.2) = 0.83, *p* = .53, 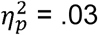, GG]. Taken with numerically weak category representations across ROIs as depicted in Figure 6A, these results show little evidence that items belonging to the same category were represented more similarly than equally similar items from different categories during observational study runs.

**Figure 6.**
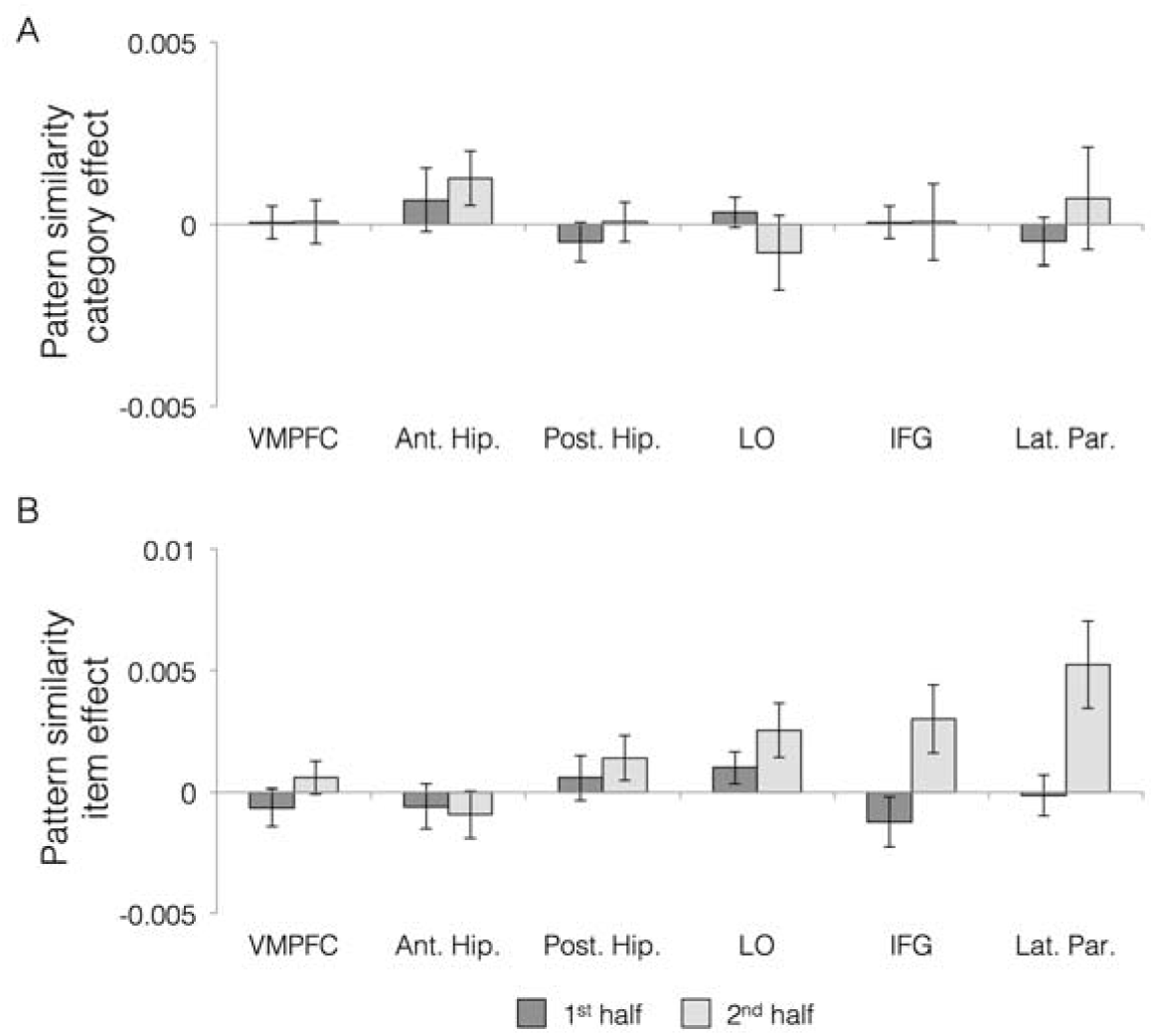
Pattern similarity analyses from observational study runs separated by first and second half of the learning phase. **A**. Category effects were defined as neural pattern similarity for items from the same category – neural pattern similarity for items from different categories, using only items that differed by four features from one another. **B**. Item effects (pattern similarity for comparisons of the same item - comparisons of items from the same category). VMPFC = ventromedial prefrontal cortex, ant. hip. = anterior hippocampus, post. hip. = posterior hippocampus, LO = lateral occipital cortex, IFG = inferior frontal gyrus, Lat. Par. = lateral parietal cortex.

In contrast, comparisons of item-level indices across ROIs and learning halves (Figure 6B) showed a significant main effect of ROI [*F*(2.9,82.4) = 3.30, *p* = .03, 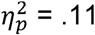, GG], a significant main effect of learning half [*F*(1,28) = 5.42, *p* = .03, 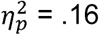], and a significant ROI x learning half interaction [*F*(3.1,86.3) = 2.79, *p* = .04, 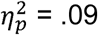, GG]. We first followed up on these significant effects by comparing the indices of item-level representations to zero to determine whether individual regions showed reliable item-level representations. There were no significant item-level representations during the first half of learning (all t’s < 1.5, p’s > .15). Reliable item-level representations emerged in the second half of training in lateral occipital cortex, inferior frontal gyrus, and lateral parietal cortex (all t’s > 2.1, p’s < .05), with no other region reaching significance (all t’s < 1.5, p’s > .15). Although the strength of item-representations was generally stronger in the second compared to first half of learning across ROIs, this effect was only reliable within individual ROIs for the inferior frontal gyrus and lateral parietal cortex (both t’s > 2.4, p’s < .03; all other t’s < 1.2, p’s > .28). Thus, we found that item-level representations emerged during observational study in several regions we predicted would represent individual category members (lateral occipital cortex, inferior frontal gyrus, and lateral parietal cortex), and that the strength of those representations increased with learning.

## Discussion

In the present study, we tested whether exemplar- and prototype-based category representations could co-exist in the brain within a single task under conditions that favor both exemplar memory and prototype extraction. We found signatures of both types of representations across distinct brain regions when participants categorized items during the learning phase. Consistent with predictions based on prior studies, the ventromedial prefrontal cortex and anterior hippocampus tracked abstract prototype information, and the inferior frontal gyrus and lateral parietal cortex tracked specific exemplar information. That inferior frontal gyrus and lateral parietal cortex represent individual items was also apparent when measured via pattern similarity analyses computed during observational study runs. In addition, we tested whether individuals relied on different types of representations over the course of learning. We did not find evidence of representational shifts either from specific to abstract or vice versa. Instead, results suggested that both types of representations emerged together during learning. Together, we show that specific and abstract representations need not form at the expense of one another and may instead exist in parallel for the same categories.

A great deal of prior work in the domain of category learning has focused on whether classification of novel category members relies on retrieval of individual category exemplars (Kruschke, 1992; Medin & Schaffer, 1978; Nosofsky, 1986) or instead on abstract category prototypes (Homa et al., 1973; Posner & Keele, 1968; Reed, 1972). These two representations are often pitted against one another with one declared the winner over the other, which is based largely on typical model-fitting procedures for behavioral data. Indeed, fitting exemplar and prototype models to behavioral data in the present study generally showed better fit of the prototype model over the exemplar model. However, using neuroimaging allowed us to detect both types of representations apparent across different parts of the brain. These results thus contribute to the ongoing debate about the nature of category representations in behavioral studies of categorization by showing that individuals may maintain multiple representations simultaneously even when one model shows better overall fit to behavior.

In addition to contributing novel findings to a longstanding debate in the behavioral literature, the present study also helps to resolve between prior neuroimaging studies fitting prototype and exemplar models to brain data. Specifically, two prior studies found conflicting results: one study found only exemplar representations in the brain (Mack et al., 2013) whereas another found only prototype representations (Bowman & Zeithamova, 2018). Notably the brain regions tracking exemplar predictions were different than those identified as tracking prototype predictions, showing that these studies engaged different brain systems in addition to implicating different categorization strategies. However, because the category structures, stimuli and analysis details also differed between these studies, the between-studies differences in the identified neural systems could not be uniquely attributed to the distinct category representations that participants presumably relied on. The present data newly show that neural prototype and exemplar correlates can exist not only across different task contexts but also within the same task, providing strong evidence that these neural differences reflect distinct category representations rather than different task details. Moreover, our results aligned with those found separately across two studies, replicating the role of the VMPFC and anterior hippocampus in tracking prototype information (Bowman & Zeithamova, 2018) and replicating the role of inferior prefrontal and lateral parietal cortices in tracking exemplar information (Mack et al., 2013).

What causes differences in the types of representations engaged across tasks and why were we able to detect both in the present study? A number of manipulations are known to influence which memory systems are engaged to support categorization, including whether training items are relatively coherent or more variable (Bowman and Zeithamova, in press), whether training is intentional or incidental (Aizenstein et al., 2000; Reber, Gitelman, Parrish, & Mesulam, 2003), whether individuals learn single or contrasting categories (Zeithamova et al., 2008), and whether a category member follows a category rule or is a rule-breaking exception (Davis, Love, & Preston, 2012). In designing the present study, we included a relatively coherent category structure that was likely to promote prototype formation (Bowman and Zeithamova, in press, 2018), but balanced it with an observational rather than feedback-based training task in hopes of emphasizing individual items and promoting exemplar representations. Detecting exemplar-tracking regions using this design provides new evidence that such observational training can enhance formation of exemplar representations relative to feedback-based training, consistent with evidence that these two forms of training engage different neural systems (Cincotta & Seger, 2007; Shohamy et al., 2004).

While co-existing prototype and exemplar representations were clear during the learning phase of our task, they were less clear during the final test phase. The VMPFC and anterior hippocampus continued to track prototypes during the final test, but exemplar-tracking regions no longer emerged. These weaker exemplar correlates in the brain were also matched by a weaker exemplar effect in behavior. While we observed a reliable advantage for old relative to new items matched for distance during the interim tests, the old advantage was no longer significant in the final test. The effect size for the prototype advantage in model fits was also larger in the final test than in the learning phase. This finding was unexpected, but we offer several possibilities that can be investigated further in future research. One possibility is that exemplar representations were weakened in the absence of further observational study runs that had boosted exemplars in earlier phases. Similarly, framing it as a ‘final test’ may have switched participants from trying to gather multiple kinds of information that might improve later performance (i.e., both exemplar and prototype) to simply deploying the strongest representation that they had, which seems to have been prototype-based. Alternatively, there may be real, non-linear dynamics in how prototype and exemplar representations develop. For example, exemplar representations may increase up to some threshold while individuals are encoding these complex stimuli, then decrease as a result of repetition suppression (Desimone, 1996; Gonsalves, Kahn, Curran, Norman, & Wagner, 2005; Henson, Shallice, Gorno-Tempini, & Dolan, 2002) once individual items are sufficiently well represented. Of course, future studies will be needed to both replicate this finding and directly test these differing possibilities.

In addition to measuring neural exemplar and prototype correlates during the categorization tests, we used pattern similarity analyses as a complementary means of assessing item-level and category-level information during the observational study runs. We found that exemplar-tracking regions, as measured with model-based MRI during generalization (IFG and lateral parietal cortex), also showed representations of individual items during the study runs, as measured by pattern similarity. Moreover, exemplar correlates and item representations were defined from separate data, providing independent confirmation that IFG and lateral parietal cortex contain representations of individual category members and that these representations can be successfully indexed using either method. These results are consistent not only with prior work showing exemplar correlates in these regions during categorization (Mack et al., 2013), but also with the larger literature on their role in maintaining memory specificity. In particular, IFG is thought to play a critical role in resolving interference between similar items (Badre & Wagner, 2005; Bowman & Dennis, 2016; Jonides, Smith, Marshuetz, Koeppe, & Reuter-Lorenz, 1998; Kuhl, Dudukovic, Kahn, & Wagner, 2007) while lateral parietal cortices often show high fidelity representations of individual items and features necessary for task performance (Kuhl & Chun, 2014; Xiao et al., 2017). The present findings support and further this prior work by showing that regions supporting memory specificity across many memory tasks also contribute to both learning and generalization of new categories.

Unlike item-level representations, we did not find evidence for category-level representations as measured by neural pattern similarity. The lack of category-level representations may appear to conflict with model-based MRI results showing both prototype- and exemplar-based category representations, but differences in these approaches may be meaningful. First, as mentioned above, the model-based MRI analyses and pattern similarity analyses were computed from separate data, generalization tests and observational study, respectively. It is possible that different demands during these two phases made it difficult to detect category representations during observational study that were more apparent during categorization. Observational study may have pushed participants toward encoding item-level information and focusing on the unique combinations of features that differentiated one training item from other similar ones. Furthermore, it is possible that abstract representations may be formed on the fly in response to generalization demands (Banino, Koster, Hassabis, & Kumaran, 2016; Carpenter & Schacter, 2017, 2018). Such generalization demands would be present at categorization tests but not during observational learning. Alternatively, prior work has shown that category learning can cause items within a category to be perceived as more similar to one another and/or items from different categories to be perceived as less similar to one another following learning (Beale & Keil, 1995; Goldstone, 1994; Livingston, Andrews, & Harnad, 1998). These categorical perception effects are thought to be driven by biases in attention toward relevant features and/or away from irrelevant features (Goldstone & Steyvers, 2001; Medin & Schaffer, 1978; Nosofsky, 1986), and can change neural representations of category members (Folstein, Palmeri, & Gauthier, 2013; Freedman, Riesenhuber, Poggio, & Miller, 2001; Myers & Swan, 2012). However, our categories did not have separate relevant and irrelevant features, as all features were equally relevant. It has been unclear from prior work whether these perceptual shifts occur when the category structure does not drive attention toward some features over others. Thus, it is possible that the equal relevance of all features reduced categorical perception in our study and prevented the detection of category-level information through pattern similarity. While further studies are needed to adjudicate between these possibilities, these results indicate that model-based fMRI and pattern similarity analyses may provide complementary information about the nature of category representations.

In addition to identifying multiple, co-existing types of category representations during learning, we sought to test whether there were representational shifts as category knowledge developed. We found no evidence for a shift between exemplar and prototype representations over the course of learning. Both prototype and exemplar correlates showed numerical increases across learning in brain and behavior, suggesting strengthening of both types of representations in parallel. Prior work has suggested that there may be representational shifts during category-learning, but rather than shifting between exemplar and prototype representations, early learning may be focused on detecting simple rules and testing multiple hypotheses (Johansen & Palmeri, 2002; Nosofsky, Palmeri, & McKinley, 1994; Paniukov & Davis, 2018), whereas similarity-based representations such as prototype and exemplar representations may develop later in learning (Johansen & Palmeri, 2002). Our findings are consistent with this framework, with strong prototype and exemplar representations emerging across distinct regions primarily in the second half of learning. Our results are also consistent with recent neuroimaging studies showing multiple memory representations forming in parallel without need for competition (Collin et al., 2015; Schlichting et al., 2015), potentially allowing individuals to flexibly use prior experience based on current decision-making demands.

## Conclusion

In the present study, we found evidence for multiple types of category representations co-existing across distinct brain regions within the same categorization task. Further, the regions identified as prototype-tracking (anterior hippocampus and VMPFC) and exemplar-tracking (IFG and lateral parietal cortex) in the present study align with prior studies that have found only one or the other. These findings shed light on the multiple memory systems that contribute to concept representation and provide novel evidence of how the brain may flexibly represent information at different levels of specificity without the need for competition between representations.

## Method

### Participants

Forty volunteers were recruited from the University of Oregon and surrounding community and were financially compensated for their research participation. This sample size was determined based on effect sizes for neural prototype-tracking and exemplar-tracking regions estimated from prior studies (Bowman & Zeithamova, 2018; Mack et al., 2013), allowing for detection of these effects with at least 80% power. All participants provided written informed consent, and Research Compliance Services at the University of Oregon approved all experimental procedures. All participants were right-handed, native English speakers and were screened for neurological conditions, medications known to affect brain function, and contraindications for MRI.

A total of 11 subjects were excluded: 3 subjects for chance performance (< .6 by the end of the training phase and/or < .6 for trained items in the final test), 5 subjects for high correlation between regressors (r > .9 for prototype and exemplar regressors and/or rank deficiency across the entire design), 1 subject for excessive motion (> 1.5 mm within multiple runs), and 2 subjects for failure to complete all phases. This left 29 subjects (age: *M* = 21.9 years, *SD* = 3.3 years, range 18-30 years; 19 females) reported in all analyses. Additionally, we excluded single runs from three subjects who had excessive motion limited to that single run.

### Materials

Stimuli consisted of cartoon animals that differed on eight binary features: neck (short vs. long), tail (straight vs. curled), foot shape (claws vs. round), snout (rounded vs. pig), head (ears vs. antennae), color (purple vs. red), body shape (angled vs. round), and design on the body (polka dots vs. stripes) (Bozoki et al., 2006; Zeithamova et al., 2008; available for download osf.io/8bph2). The two possible versions of all features can be seen across the two prototypes shown in Figure 1C. For each participant, the stimulus that served as the prototype of category A was randomly selected from four possible stimuli and all other stimuli were re-coded in reference to that prototype. The stimulus that shared no features with the category A prototype served as the category B prototype. Physical distance between any pair of stimuli was defined by their number of differing features. Category A stimuli were those that shared more features with the category A prototype than the category B prototype. Category B stimuli were those that shared more features with the category B prototype than the category A prototype. Stimuli equidistant from the two prototypes were not used in the study.

### Training set

The training set included four stimuli per category, each differing from their category prototype by two features (see Table 2 for training set structure). The general structure of the training set with regard to the category prototypes was the same across subjects, but the exact stimuli differed based on the prototypes selected for a given participant. The training set structure was selected to generate many pairs of training items that were four features apart both within the same category and across the two categories. This design ensured that categories 1) could not be learned via unsupervised clustering based on similarity of exemplars alone and 2) to control the number of shared features between items when computing pattern similarity within and between categories, equating for physical similarity when determining the effect of category membership.

**Table 2.** Dimension values for example prototypes and training stimuli from each category.

|  | Dimension values |  |  |  |  |  |  |  |
| --- | --- | --- | --- | --- | --- | --- | --- | --- |
| Stimulus | <u>1</u> | <u>2</u> | <u>3</u> | <u>4</u> | <u>5</u> | <u>6</u> | <u>7</u> | <u>8</u> |
| Prototype A | 1 | 1 | 1 | 1 | 1 | 1 | 1 | 1 |
| A1 | 1 | 1 | 1 | 1 | 1 | 1 | 0 | 0 |
| A2 | 0 | 1 | 1 | 1 | 0 | 1 | 1 | 1 |
| A3 | 1 | 0 | 1 | 0 | 1 | 1 | 1 | 1 |
| A4 | 1 | 1 | 0 | 1 | 1 | 0 | 1 | 1 |
| Prototype B | 0 | 0 | 0 | 0 | 0 | 0 | 0 | 0 |
| B1 | 0 | 0 | 0 | 0 | 0 | 0 | 1 | 1 |
| B2 | 1 | 0 | 0 | 0 | 1 | 0 | 0 | 0 |
| B3 | 0 | 1 | 0 | 1 | 0 | 0 | 0 | 0 |
| B4 | 0 | 0 | 1 | 0 | 0 | 1 | 0 | 0 |

### Interim test sets

Stimuli in the interim generalization tests included 22 unique stimuli: the 8 training stimuli, both prototypes, and two new stimuli at each distance (1, 2, 3, 5, 6, 7) from the category A prototype. Distance 1, 2 and 3 items were scored as correct when participant labeled them as category A members. Items at the distance 5, 6 and 7 from the category A prototype (thus distance 3, 2, and 1 from the B prototype) were scored as correct when participant labeled them as category B members. While new unique distance 1-3, 5-7 items were selected for each interim test set, the old training stimuli and the prototypes were necessarily the same for each test.

### Final test set

Stimuli in the final test included 58 unique stimuli. Forty-eight of those consisted of 8 new stimuli selected at each distance 1-3, 5-7 from the category A prototype, each presented once during the final test. These new items were distinct from those used in either the training set or the interim test sets with the exception of the items that differed by only 1 feature from their respective prototypes. Because there are only 8 distance 1 items for each prototype, they were all used as part of the interim test sets before being used again in the final test set. The final test also included the 8 training stimuli and the two prototypes, each presented twice in this phase (Bowman & Zeithamova, 2018; Kéri, Kelemen, Benedek, & Janka, 2001; J David Smith, Redford, & Haas, 2008).

### Experimental Design

The study consisted of two sessions: one session of neuropsychological testing and one experimental session. Only results from the experimental session are reported in the present manuscript. In the experimental session, subjects underwent four cycles of observational study and interim generalization tests (Figure 1D), followed by a final generalization test (Figure 1E), all while undergoing fMRI.

In each run of observational study, participants were shown individual animals on the screen with a species label (Febbles and Badoons) and were told to try to figure out what makes some animals Febbles and others Badoons without making any overt responses. Each stimulus was presented on the screen for 5 seconds followed by a 7 second ITI. Within each study run, participants viewed the training examples three times in a random order. After two study runs, participants completed an interim generalization test. Participants were shown cartoon animals without their labels and classified them into the two species without feedback. Each test stimulus was presented for 5 seconds during which time they could make their response, followed by a 7 second ITI. After four study-test cycles, participants completed a final categorization test, split across four runs. As in the interim tests, participants were asked to categorize animals into one of two imaginary species (Febbles and Badoons) using the same button press while the stimulus was on the screen. Following the MRI session, subjects were asked about the strategies they used to learn the categories, if any, and then indicated which version of each feature they thought was most typical for each category. Lastly, subjects were verbally debriefed about the study.

### fMRI Data Acquisition

Raw MRI data are available for download via OpenNeuro (openneuro.org/datasets/ds002813). Scanning was completed on a 3T Siemens MAGNETOM Skyra scanner at the University of Oregon Lewis Center for Neuroimaging using a 32-channel head coil. Head motion was minimized using foam padding. The scanning session started with a localizer scan followed by a standard high-resolution T1-weighted MPRAGE anatomical image (TR 2500 ms; TE 3.43 ms; TI 1100 ms; flip angle 7°; matrix size 256 256; 176 contiguous slices; FOV 256 mm; slice thickness 1 mm; voxel size 1.0 1.0 1.0 mm; GRAPPA factor 2). Then, a custom anatomical T2 coronal image (TR 13,520 ms; TE 88 ms; flip angle 150°; matrix size 512 512; 65 contiguous slices oriented perpendicularly to the main axis of the hippocampus; interleaved acquisition; FOV 220 mm; voxel size 0.4 0.4 2 mm; GRAPPA factor 2) was collected. This was followed by 16 functional runs using a multiband gradient echo pulse sequence [TR 2000 ms; TE 26 ms; flip angle 90°; matrix size 100 100; 72 contiguous slices oriented 15° off the anterior commissure–posterior commissure line to reduced prefrontal signal dropout; interleaved acquisition; FOV 200 mm; voxel size 2.0 2.0 2.0 mm; generalized autocalibrating partially parallel acquisitions (GRAPPA) factor 2]. One hundred and forty-five volumes were collected for each observational study run, 133 volumes for each interim test run, and 103 volumes for each final test run.

### Statistical Analysis

#### Behavioral accuracies

##### Interim tests

To assess changes in generalization accuracy across train-test cycles, we computed a 4 (interim test run: 1-4) x 4 (distance: 0-3) repeated-measures ANOVA on accuracy for new items only. We were particularly interested in linear effects of interim test run and distance. We also tested whether there was a difference across training in accuracy for the training items themselves versus new items at the same distance from their prototypes, which can index how much participants learn about specific items above-and-beyond what would be expected based on their typicality. We thus computed a 4 (interim test run: 1-4) x 2 (item type: training, new) repeated-measures ANOVA on accuracies for distance 2 items.

##### Final test

First, to assess the effect of item typicality, classification performance in the final test (collapsed across runs) was assessed by computing a one-way, repeated-measures ANOVA across new items at distances (0-3) from either prototype. Second, we assessed whether there was an old-item advantage by comparing accuracy for training items and new items of equal distance from prototypes (distance 2) using a paired-samples t-test. For all analyses (including fMRI analyses described below), a Greenhouse-Geisser correction was applied whenever the assumption of sphericity was violated as denoted by ‘GG’ in the results.

#### Prototype and exemplar model fitting

As no responses were made during the study runs, prototype and exemplar models were only fit to test runs – interim and final tests. As the number of trials in each interim test was kept low to minimize exposure to non-training items during the learning phase, we concatenated across interim tests 1 and 2 and across interim tests 3 and 4 to obtain more robust model fit estimates for the first half vs. second half of the learning phase. Model fits for the final test were computed across all four runs combined. Each model was fit to trial-by-trial data in individual participants.

### Prototype similarity

As in prior studies (Bowman & Zeithamova, 2018; Maddox et al., 2011; Minda & Smith, 2001), the similarity of each test stimulus to each prototype was computed, assuming that perceptual similarity is an exponential decay function of physical similarity (Shepard, 1957), and taking into account potential differences in attention to individual features. Formally, similarity between the test stimulus and the prototypes was computed as follows:

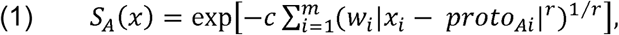

where *S*_*A*_ (*x*) is the similarity of item *x* to category A, *x*_*i*_ represents the value of the item *x* on the *i*th dimension of its *m* binary dimensions (*m* = 8 in this study), proto_A_ is the prototype of category A, *r* is the distance metric (fixed at 1 for the city-block metric for the binary dimension stimuli). Parameters that were estimated from each participant’s pattern of behavioral responses were *w* (a vector with 8 weights, one for each of the 8 stimulus features and constrained to sum to 1) and *c* (sensitivity: the rate at which similarity declines with physical distance, constrained to be 0-100).

### Exemplar similarity

Exemplar models assume that categories are represented by their individual exemplars, and that test items are classified into the category with the highest summed similarity across category exemplars (Figure 1A). As in prior studies (Nosofsky, 1987; Zaki, Nosofsky, Stanton, & Cohen, 2003), similarity of each test stimulus to the exemplars of each category was computed as follows:

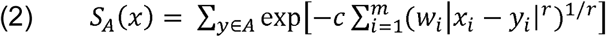

where *y* represents the individual training stimuli from category A, and the remaining notation and parameters are as in Equation 1.

### Parameter estimation

For both models, the probability of assigning a stimulus *x* to category A is equal to the similarity to category A divided by the summed similarity to categories A and B, formally, as follows:

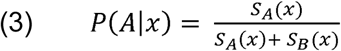

Using these equations, the best fitting *w*_1-8_ (attention to each feature) and *c* (sensitivity) parameters were estimated from the behavioral data of each participant, separately for the first half of the learning phase, second half of the learning phase, and the final test, and separately for the prototype and exemplar models. To estimate these parameters for a given model, the trial-by-trial predictions generated by Equation 3 were compared with the participant’s actual series of responses, and model parameters were tuned to minimize the difference between predicted and observed responses. An error metric (negative log likelihood of the entire string of responses) was computed for each model by summing the negative of log-transformed probabilities, and this value was minimized by adjusting *w* and *c* using standard maximum likelihood methods, implemented in MATLAB (Mathworks, Natick, MA), using the “fminsearch” function.

### Group analyses

After optimization, we computed a 2 (model: prototype, exemplar) x 2 (learning phase half: 1^st^, 2^nd^) repeated-measures ANOVA on the model fit values (i.e., negative log likelihood) to determine which model provided a better fit to behavioral responses and if there were shifts across learning in which model fit best. We used a paired-samples t-test comparing model fits during the final test to determine whether the group as a whole was better fit by the prototype or exemplar model by the end of the experiment. Finally, we used trial-by-trial similarity metrics generated by each model as regressors for the neuroimaging data (described in the fMRI analysis section) to identify brain regions tracking predictions of each model during each phase.

#### fMRI Preprocessing

The raw data were converted from dicom files to nifti files using the dcm2niix function from MRIcron (https://www.nitrc.org/projects/mricron). Functional images were skull-stripped using BET (Brain Extraction Tool), which is part of FSL (www.fmrib.ox.ac.uk/fsl). Within-run motion correction was computed using MCFLIRT in FSL to realign each volume to the middle volume of the run. Across-run motion correction was then computed using ANTS (Advanced Normalization Tools) by registering the first volume of each run to the first volume of the first functional run (i.e., the first training run). Each computed transformation was then applied to all volumes in the corresponding run. Brain-extracted and motion-corrected images from each run were entered into the FEAT (fMRI Expert Analysis Tool) in FSL for high-pass temporal filtering (100 s) and spatial smoothing (4 mm FWHM kernel for univariate analyses, 2 mm for pattern analyses).

#### Regions of interest

Regions of interest (ROIs, Figure 2) were defined anatomically in individual participants’ native space using the cortical parcellation and subcortical segmentation from Freesurfer version 6 (https://surfer.nmr.mgh.harvard.edu/) and collapsed across hemispheres to create bilateral masks. Past research has indicated that there may be a functional gradient along the hippocampal long axis, with detailed, find-grained representations in the posterior hippocampus and increasingly coarse, generalized representations proceeding toward the anterior hippocampus (Brunec et al., 2018; Frank, Bowman, & Zeithamova, 2019; Poppenk, Evensmoen, Moscovitch, & Nadel, 2013). As such, we divided the hippocampal ROI into anterior/posterior portions at the middle slice. When a participant had an odd number of hippocampal slices, the middle slice was assigned to the posterior hippocampus. Based on our prior report (Bowman & Zeithamova, 2018), we expected the anterior portion of the hippocampus to track prototype predictors, together with VMPFC (medial orbitofrontal label in Freesurfer). Based on the prior study by Mack and colleagues (2013), we expected lateral occipital cortex, inferior frontal gyrus (combination of pars opercularis, pars orbitalis, and pars triangularis freesurfer labels), and lateral parietal cortex (combination of inferior parietal and superior parietal freesurfer labels) to track exemplar predictors. The posterior hippocampus was also included as an ROI, to test for an anterior/posterior dissociation within the hippocampus. While one might expect the posterior hippocampus to track exemplar predictors based on the aforementioned functional gradient, our prior report (Bowman and Zeithamova, 2018) found only a numeric trend in this direction and Mack et al. (2013) did not report any hippocampal findings despite significant exemplar correlates found in the cortex. Thus, we did not have strong predictions regarding the posterior hippocampus, other than being distinct from the anterior hippocampus.

#### Model-based fMRI analyses

fMRI data were modeled using the GLM. Three task-based regressors were included in the GLM: one for all trial onsets, one that included modulation for each trial by prototype model predictions, and one that included modulation for each trial by exemplar model predictions. The regressor for all trial onsets was included to account for activation that is associated with performing a categorization task generally, but does not track either model specifically. The modulation values for each model were computed as the summed similarity across category A and category B (denominator of Equation 3) generated by the assumptions of each model (from Equations 1 and 2). This summed similarity metric indexes how similar the current item is to the existing category representations as a whole (regardless of which category it is closer to) and has been used by prior studies to identify regions that contain such category representations (Bowman & Zeithamova, 2018; Davis & Poldrack, 2014; Mack et al., 2013). By including both model predictors in the same GLM, we account for any shared variance between the regressors. Events were modeled with a duration of 5 seconds, which was the fixed length of the stimulus presentation. Onsets were then convolved with the canonical hemodynamic response function as implemented in in FSL (a gamma function with a phase of 0 seconds, and SD of 3 seconds, and a mean lag time of 6 seconds). Finally, the six standard timepoint-by-timepoint motion parameters were included as regressors of no interest.

For region of interest analyses, we obtained an estimate of how much the BOLD signal in each region tracked each model predictor by dividing the mean ROI parameter estimate by the standard deviation of parameter estimates (i.e., computing an effect size measure). Normalizing the beta values by their error of the estimate de-weighs values associated with large uncertainty, similar to how lower level estimates are used in group analyses as implemented in FSL (S. M. Smith et al., 2004). These normalized beta values were then averaged across the appropriate runs (interim tests 1-2, interim tests 3-4, all four runs of the final test) and submitted to group analyses.

We tested whether prototype and exemplar correlates emerge across different regions and/or at different points during the learning phase. To do so, we computed a 2 (model: prototype, exemplar) x 2 (learning phase: 1^st^ half, 2^nd^ half) x 6 (ROI: VMPFC, anterior hippocampus, posterior hippocampus, lateral occipital, lateral prefrontal, and lateral parietal cortices) repeated-measures ANOVA on parameter estimates from the interim test runs. We were interested in a potential model x ROI interaction effect, indicating differences across brain regions in the type of category information represented. Following any significant interaction effect, we computed one-sample t-tests to determine whether each region significantly tracked a given model. Given *a priori* expectations about the nature of these effects, we computed one-tailed tests only on the effects of interest: for example, in hypothesized prototype-tracking ROIs (anterior hippocampus and VMPFC), we computed one-sample t-tests to compare prototype effects to zero. We followed a similar procedure in hypothesized exemplar-tracking ROIs (inferior frontal gyrus, lateral parietal cortex, lateral occipital cortex). We were also interested in potential interactions with the learning phase, which would indicate shift across learning in category representations. Following any such interaction, follow-up ANOVAs or t-tests were performed to better understand drivers of the effect.

We next tested ROI differences in the final generalization phase. To do so, we computed a 2 (model: prototype, exemplar) x 6 (ROI: see above) repeated-measures ANOVA on parameter estimates from the final generalization test. We were particularly interested in the model x ROI interaction effect, which would indicate that regions differ in which model they tracked.

#### Pattern similarity analyses during observational study runs

As a complementary analysis, we also examined item-level and category-level representations using pattern similarity analyses during observational study runs. Study runs contained many pairs of items, both from the same category and from opposing categories, that were equated on physical similarity (four features apart). To generate beta-series representing a pattern of activation for each trial, a GLM was run in which each trial was modeled as a separate regressor of interest. In addition to the 24 regressors for the trials (8 category examples repeated 3 times), the model included 6 standard motion regressors, totaling 30 regressors for each run. As with the model-based analyses described above, the duration was set to 5 seconds and regressors were convolved with the canonical hemodynamic response function. After estimating the model for each run, we concatenated the single-trial betas across all training runs.

Following concatenation, we submitted each beta-series to pattern similarity analyses using the PyMVPA toolbox (www.pymvpa.org) for Python (www.python.org). For each ROI, we extracted the vector of beta parameters across voxels for two trials of interest and correlated them using a Pearson’s correlation. Within each half of training (study runs 1-4 vs. study runs 5-8), we computed the pattern similarity score for all trials not in the same run (all run 1 trials compared to all run 2, 3, and 4 trials; all run 2 trials with all run 3 and 4 trials, etc.). Pattern similarity was never computed within a run to avoid issues with local autocorrelations (Mumford et al., 2014). After computing all similarity scores, the resulting r-values were Fisher z-transformed. Three types of similarity scores of interest were then extracted: same item comparisons, same category comparisons, and different category comparisons. For same item comparisons, the similarity score represents the correlation in activation patterns when the exact same item is repeated. For same category comparisons, the similarity score represents the correlation in activation patterns for two items that are in the same category, but are not the exact same item. For different category comparisons, the similarity score represents the correlation in activation patterns for two items that are from opposing categories. To ensure that differences in similarity scores between items in the same vs. different categories was not due only to differences in physical similarity, we only extracted similarity scores in which items were 4 features different from one another for both the same category and different category condition.

To measure item-level representations, we then subtracted the same category similarity from same item similarity (same item – same category). To measure category-level representations, we subtracted the different category similarity from the same category similarity (same category – different category). To test for differences in the strength of item and category information across ROIs and across learning, we submitted these differences scores to separate 2 (learning phase) x 6 (ROI) repeated-measures ANOVAs. Following a significant effect of ROI or ROI x learning phase interaction, we computed one-sample t-tests within each ROI to determine whether the pattern similarity effect of interest was present (i.e., item or category representation significantly different than zero). Following a significant interaction, we used paired-samples t-tests to identify differences between the first and second half of learning within each ROI.

## Acknowledgements

Funding for this work was provided in part by the Lewis Family Endowment, which supports the Robert and Beverly Lewis Center for Neuroimaging at the University of Oregon (D.Z.), by the National Institute on Aging Grant F32-AG054204 (C.R.B).

## Notes

### Competing Interest Statement

The authors have declared no competing interest.

### Summary of Updates

Correction to title

